# Membrane-Tethered Mucin 1 is Stimulated by Interferon in Multiple Cell Types and Antagonizes Influenza A Virus Infection in Human Airway Epithelium

**DOI:** 10.1101/2021.03.11.434997

**Authors:** Ethan Iverson, Kira Griswold, Daniel Song, Talita B. Gagliardi, Kajal Hamidzadeh, Mehmet Kesimer, Sanju Sinha, Melissa Perry, Gregg A. Duncan, Margaret A. Scull

**Affiliations:** Department of Cell Biology & Molecular Genetics, Maryland Pathogen Research Institute, University of Maryland, College Park, MD, USA; Fischell Department of Bioengineering, University of Maryland, College Park, MD, USA; Marsico Lung Institute, University of North Carolina, Chapel Hill, NC, USA; Cancer Data Science Laboratory, National Cancer Institute, National Institutes of Health, Bethesda, MD, USA; Center for Bioinformatics and Computational Biology, University of Maryland, College Park, MD, USA

**Keywords:** influenza virus, mucin, interferon, airway epithelium, macrophage

## Abstract

Influenza A virus (IAV) causes seasonal epidemics and periodic pandemics, resulting in significant morbidity and mortality in the human population. Tethered mucin 1 (MUC1) is highly expressed in airway epithelium, the primary site of IAV replication, and also by other cell types that influence IAV infection, including macrophages. MUC1 has the potential to influence infection dynamics through physical interactions and/or signaling activity, and recent work suggests MUC1 acts as a releasable decoy receptor and anti-inflammatory molecule during IAV infection. Still, the modulation of MUC1 and its impact during viral pathogenesis remains unclear. Thus, we sought to further investigate the interplay between MUC1 and IAV in an *in vitro* model of primary human airway epithelium (HAE). Our data indicate that a recombinant IAV hemagglutinin (H3) and H3N2 virus can bind endogenous HAE MUC1. We find that infection of HAE cultures with H1N1 or H3N2 IAV strains does not trigger enhanced MUC1 shedding, but instead stimulates an increase in cell-associated MUC1 protein. We observed a similar increase after stimulation with either type I or type III interferon (IFN); however, inhibition of IFN signaling during H1N1 infection only partially abrogated this increase, indicating multiple soluble factors contribute to MUC1 upregulation during the antiviral response. We expanded these findings and demonstrate that in addition to HAE, primary human monocyte-derived macrophages also upregulate MUC1 protein in response to both IFN treatment and conditioned media from IAV-infected HAE cultures. We then developed HAE genetically depleted for MUC1 to determine its impact on IAV pathogenesis, finding that MUC1 knock-out cultures exhibited enhanced viral growth compared to control cultures for several IAV strains. Together, our data support a model whereby MUC1 antagonizes productive uptake of IAV in HAE. Infection then stimulates MUC1 expression on multiple cell types through IFN-dependent and -independent mechanisms that may further impact infection dynamics.

**Author Summary:** The mucosal surface of the respiratory epithelium is an important site of first contact for viral respiratory pathogens. Large and heavily glycosylated molecules known as tethered mucins extend from the cell surface and may physically restrict access to underlying cells. Recently, one of these tethered mucins, MUC1, has also been shown to influence cell signaling and inflammation. Still, despite its abundance in the airway and multifunctional capability, the role of MUC1 during influenza virus infection in the human respiratory tract remains unclear. Here, we demonstrate that influenza virus directly interacts with MUC1 in a physiologically-relevant model of human airway epithelium and find that MUC1 protein expression is elevated throughout the epithelium and in primary human monocyte-derived macrophages in response to important antiviral signals produced during infection. Using genetically-modified human airway cultures lacking MUC1, we then provide evidence of more efficient influenza virus infection in the absence of this mucin. Our data suggest that MUC1 not only physically restricts influenza virus uptake, but also represents a dynamic component of the host response that acts to further stem viral spread.

## Introduction

The respiratory epithelium encodes large and extensively glycosylated proteins, termed mucins, to maintain airway surface hydration and protect the underlying cells from environmental insults, such as respiratory viruses [1,2]. While some mucins are secreted and form a mucus gel, others – the aptly named “tethered” mucins – remain anchored to the apical epithelial cell surface, giving rise to the periciliary layer (PCL) [1–3]. The PCL serves as a platform for overlying secreted mucins, allowing ciliary action to propel the secreted mucus gel in a process known as mucociliary clearance (MCC) [4,5]. Additionally, tethered mucins of the PCL represent steric obstacles to frustrate further access to the underlying epithelium [2]. In addition to the bulky extracellular domain (ED) typical of tethered mucins, the highly abundant mucin 1 (MUC1) features a highly-conserved cytoplasmic tail (CT) that can be differentially phosphorylated [6,7] and interact with many partners including kinases and adapter proteins involved in signal transduction [3,8,9]. The presence of an autoproteolytic SEA domain upstream the transmembrane domain, in conjunction with enzymatic sheddases, can lead to the release of the MUC1-ED domain from the MUC1-CT domain [3,10]. MUC1-CT can also be translocated to the nucleus [11–13], altogether supporting important functions outside its canonical representation among the PCL.

MUC1/Muc1 (humans/mice) has been implicated in various aspects of both bacterial and viral infections. For example, the genetic disruption of *Muc1* is associated with elevated inflammation and faster *Pseudomonas aeruginosa* clearance [8], yet results in more severe *Streptococcus pneumoniae* infection [14]. Adenoviral infection in *Muc1*^*-/-*^ mice is modestly increased with no significant inflammatory differences in the lung [15] and adenoviral vector gene transfer efficiency *in vitro* and *in vivo* is inhibited by MUC1/Muc1 expression [16,17], suggesting that MUC1 restricts adenovirus by acting as a physical barrier. Outside the airway, MUC1 has been shown to be an attachment factor for *Helicobacter pylori* [18] and *Salmonella enterica* [19], while the presence of MUC1 in breast milk is protective against human immunodeficiency virus transmission [20]. MUC1 has also been shown to suppress respiratory syncytial virus-induced inflammation *in vitro* by forming a negative feedback loop with tumor necrosis factor (TNFα) [21] and altered expression of MUC1 has been described in response to multiple inflammatory stimuli [22], suggesting it might play a universal and dynamic role during insult by different pathogens [23,24]. Notably, no consensus on MUC1 function or dynamics during infection is reflected in these studies.

Influenza A virus (IAV) infects the human airway epithelium (HAE) [25] and causes an estimated annual burden of 290,000-645,000 deaths worldwide in non-pandemic years [26]. To gain access to airway epithelial cells, IAV must first penetrate the secreted mucus and underlying PCL barriers. Subsequent endocytic uptake into epithelial cells is mediated through interactions between the viral attachment protein hemagglutinin and glycans with terminal sialic acid (SA) linkages on the cell surface [27]. While it is known that SA recognition heavily impacts cellular tropism and epizootic potential [28], the extent and consequence of IAV attachment to SA on specific host proteins is unclear [29]. A recent report suggests that IAV can interact with the extracellular domain of MUC1 and that this interaction has important implications for pathogenesis *in vivo* [30]. However, it is not known if MUC1 can restrict IAV access to well-differentiated epithelial cells, or if SA-mediated interactions subvert a normally protective physical role and instead support IAV uptake. Additionally, it is not known how MUC1 expression is impacted during IAV infection of the respiratory epithelium and whether its immunomodulatory role is important in the context of IAV pathogenesis.

Here we investigate specific interactions between IAV and MUC1 in a physiologically-relevant model of HAE. Consistent with previous reports in cell lines [30], we show that IAV can interact with membrane-tethered MUC1 in HAE; however, in contrast to earlier findings, we find no evidence of IAV-mediated MUC1 shedding in several epithelial model systems. Our data instead indicate that MUC1 is upregulated in all HAE component cell types as well as primary human monocyte-derived (PMD) macrophages by soluble factors, including type I and type III interferons, produced during IAV infection. Then, using an *in vitro* HAE model system that is genetically deleted for MUC1, we demonstrate that depletion of MUC1 is pro-viral for several IAV strains, leading to enhanced IAV replication and spread.

## Results

### The IAV hemagglutinin protein binds MUC1 isolated from HAE apical secretions and co-localizes with MUC1 during infection

Previous work suggests that IAV can interact with MUC1 based on fluorescence microscopy and colocalization analysis in A549 cells [30]. Thus, we initially sought to determine if the IAV hemagglutinin protein binds MUC1 derived from an *in vitro* model of primary HAE since this system recapitulates important aspects of airway epithelial morphology and physiology including both secreted and tethered mucin expression (**S1 Fig**) [2,31,32]. Notably, in HAE cultures, MUC1 can also be identified in apical secretions along with other, less abundant tethered mucins (e.g., MUC4, MUC16) [2,33].

MUC1 enrichment from HAE secretions was achieved by immunoprecipitation with anti-MUC1-coated beads (**Fig 1A**). This MUC1-enriched material was then washed and eluted off the beads before being separated by agarose electrophoresis and transferred to a membrane, where interaction with influenza virus hemagglutinin protein was determined using a recombinant, Fc-tagged H3 hemagglutinin (rH3-Fc). Detection of rH3-Fc binding and anti-MUC1-ED reactivity in the same region of the membrane indicated a likely interaction between the viral attachment protein and this mucin molecule (**Fig 1B**). MUC16, another tethered mucin that was also previously identified in HAE secretions, was detected in the total input but not in the immunoprecipitated conditions. These data support that MUC1 was further enriched from other tethered mucins and are consistent with the conclusion that detection of rH3-Fc is indicative of hemagglutinin-MUC1 binding.

**Fig 1.**
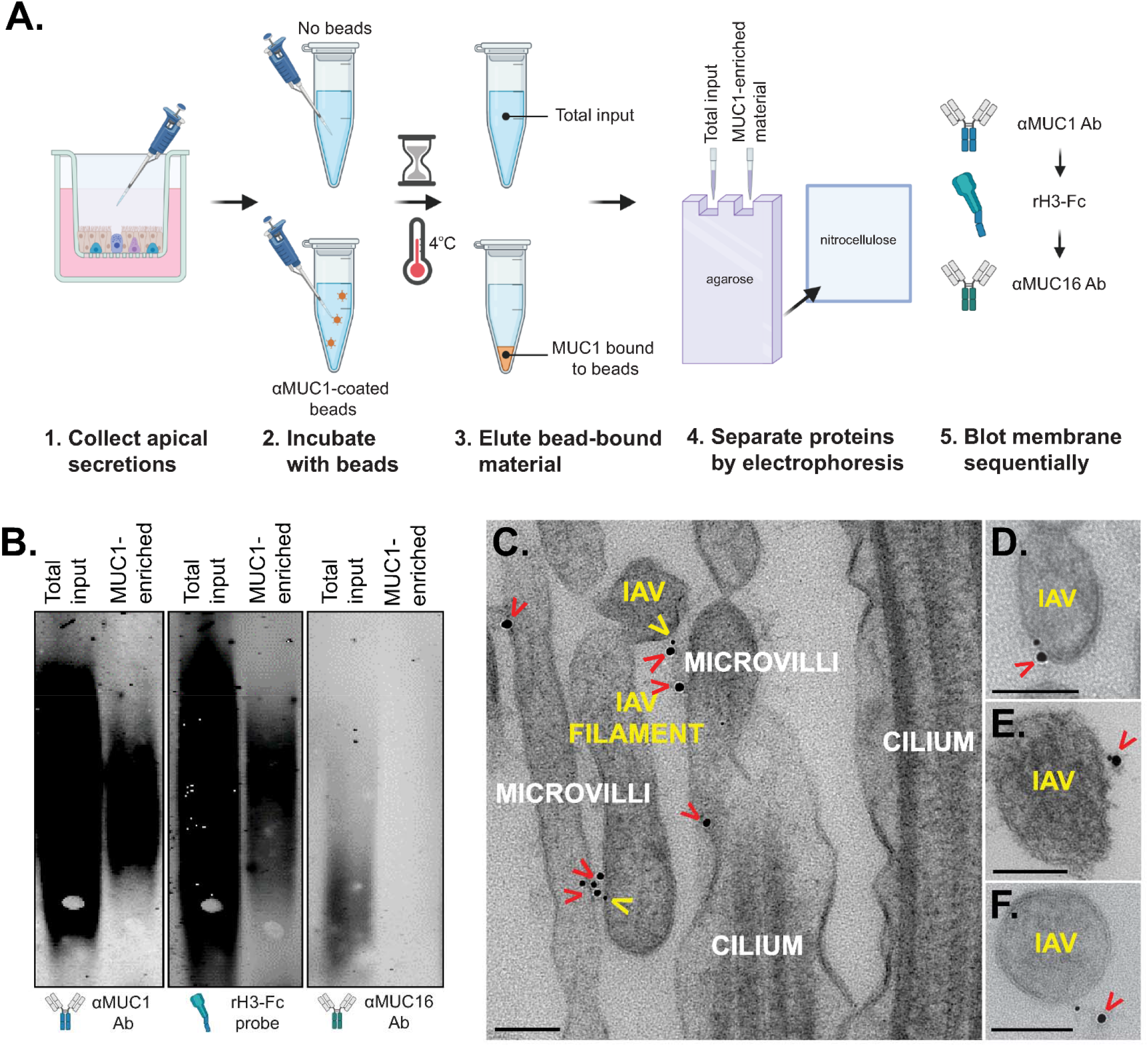
IAV hemagglutinin protein binds HAE-derived MUC1 and co-localizes with MUC1 during infection. (A) Overview of the experimental workflow used in panel B. Briefly, HAE apical secretions were collected and incubated with beads to immunoprecipitate MUC1-containing material. MUC1-enriched material was then recovered from the beads and separated (in parallel with total input material) by agarose gel electrophoresis before transfer to a membrane. (B) Samples processed as described in (A) were probed sequentially with αMUC1-ED and αMUC16 antibodies and a rH3-Fc probe. Blotting represents the same two lanes imaged with indicated probe. (C-E) Transmission electron microscopy of HAE after adsorption with A/Udorn/307/72(H3N2) influenza virus. Images were taken from (C, D, and F) primary HAE and (E) and BCi-NS1.1-derived HAE [47]. MUC1 (indicated by red carets) and H3 (yellow carets) were detected with 18nm and 6nm gold nanoparticle-conjugated antibodies, respectively. Scale bars = 100 nm. Diagram in (A) was created with BioRender.com.

To determine if the hemagglutinin-MUC1 interaction occurs in the context of the native HAE microenvironment, we utilized A/Udorn/307/72, a well-characterized strain that natively possesses an H3 similar to the recombinant H3 probe. HAE cultures were inoculated with >5 × 10^5^ plaque forming units (PFU) of virus and subsequently chilled to 4°C so as to irreversibly stabilize virus adsorption and restrict cellular entry [34]. Next, we performed transmission electron microscopy with immunogold labeling to detect IAV H3 as well as MUC1-ED, allowing us to observe potential colocalization of these two molecules prior to cellular uptake. Consistent with previous reporting, we identified MUC1 at the apical surface primarily localized to microvilli [2]. IAV was also frequently in close proximity to immunogold-labelled MUC1 (**Fig 1C-F**), in line with our *in vitro* interaction and prior work in A549 cells [30]. Taken together, our results suggest that influenza virus interacts with MUC1 during the early stages of infection in a physiologically-relevant system that recapitulates the extracellular environment in the airway.

### IAV replication in HAE is not associated with an increase in soluble MUC1

Given our results indicating HAE-MUC1 interacts with IAV hemagglutinin, we next sought to determine the consequence of this interaction. Previous work in CHO cells suggested that the ectodomain of ectopically-expressed MUC1 could act as a releasable decoy that is shed upon IAV binding to prevent subsequent infection of underlying cells [30]. To determine whether MUC1 is shed during viral challenge in the context of the airway PCL, we inoculated primary, well-differentiated HAE cultures with either A/Udorn/307/72 or another well-characterized virus, A/PR/8/34, which possesses a H1 hemagglutinin, and quantified MUC1 and infectious virus in apical washes 24 hours post-infection (hpi; **Fig 2A** and **2B**). Surprisingly, in contrast to previous observations, we found soluble MUC1 levels were reduced relative to mock-infected cultures. Both viruses resulted in similar reductions of soluble MUC1 – although neither reduction reached significance – and achieved similar viral titers with A/Udorn/307/72 being more variable than A/PR/8/34 in both of these regards.

**Fig 2.**
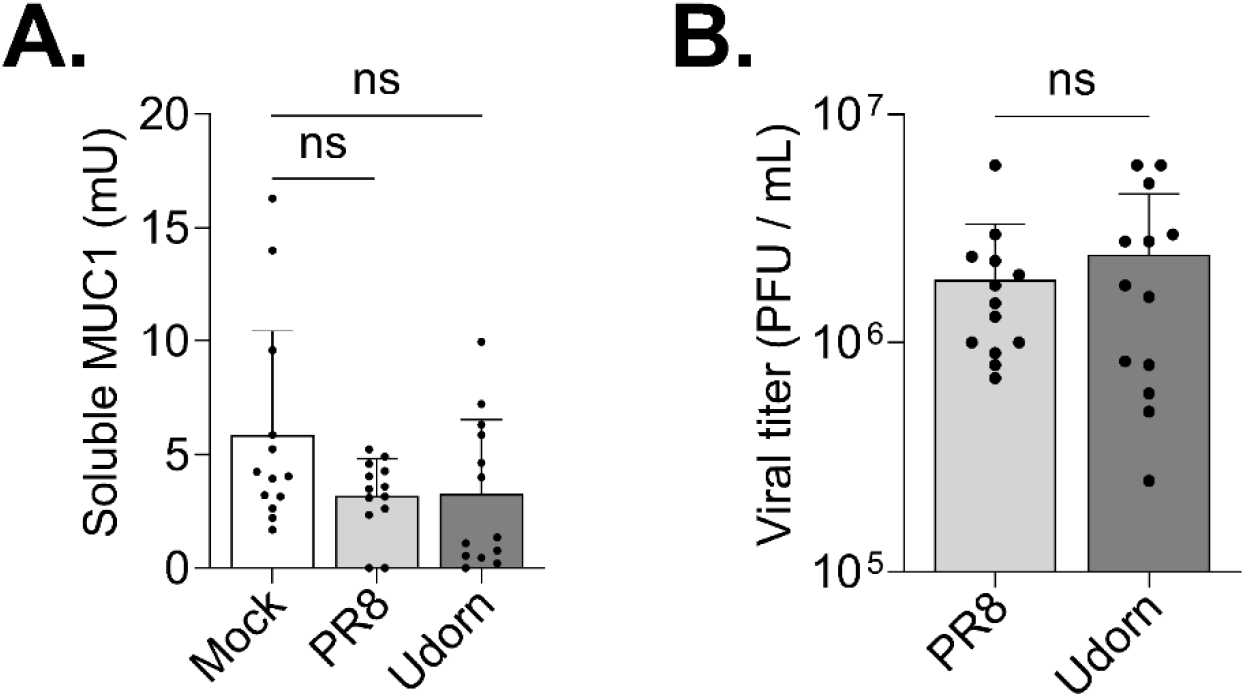
IAV replication in HAE is associated with a reduction in soluble MUC1. HAE cultures were infected with 5 × 10^4^ PFU of either A/PR/8/34 or A/Udorn/307/72 or mock-infected. After 24 hours, apical HAE compartments were washed with PBS which was used to determine (A) soluble MUC1-ED by ELISA and (B) viral titer by plaque assay. Results shown are from four independent experiments, each performed in HAE cultures derived from a unique donor. Experimental results were analyzed by Mann-Whitney U test compared to (A) mock conditions or (B) each other (ns = not significant).

To determine if a lack of MUC1 shedding after IAV challenge was an HAE-specific phenomenon, we executed a similar experiment in A549 cells expressing endogenous MUC1.Following a one hour incubation at 4°C to allow viral particles to bind to the cell surface, we removed the inoculum, returned the cultures to 37°C, and quantified MUC1 and infectious virus in cell culture supernatants 24 hours later (**S2 Fig**). Similar to our HAE results, infection with A/PR/8/34 and A/Udorn/307/72 yielded a decrease in soluble MUC1 which was significant only for A/Udorn/307/72. These data corroborate our results in HAE and together suggest that MUC1 expressed endogenously in human airway cells is not shed during IAV challenge.

### Cell-associated MUC1 levels are upregulated during IAV infection and after interferon treatment

As the reduction of soluble MUC1 levels following infection of human airway cells was unexpected, we sought to further characterize MUC1 dynamics in HAE after IAV challenge. Since previous reports have described an increase in MUC1 protein following IFNγ exposure in other systems [35], and IAV infection of HAE triggers both type I and type III IFN [36], we quantified MUC1 gene expression and cell-associated MUC1 protein levels following IAV infection, or after treatment of HAE with IFNβ, IFNλ3, or TNFα (previously implicated in upregulating MUC1 [23,37]). Neither type I or type III IFN treatment (**Fig 3A** and **3B**) nor IAV infection (**Fig 3C**) triggered an increase in MUC1 transcripts above mock-treated controls, let alone a response typical of canonical interferon stimulated genes (**S3 Fig**). However, type I and type III IFN, along with IAV, were able to stimulate production of MUC1 protein similar to that seen with TNFα (**Fig 3D**). Furthermore, IAV-mediated upregulation of MUC1 protein was at least partially IFN signaling-independent, as the addition of a Janus tyrosine kinase (JAK)1 inhibitor did not abolish this increased protein expression (**Fig 3E**).

**Fig 3.**
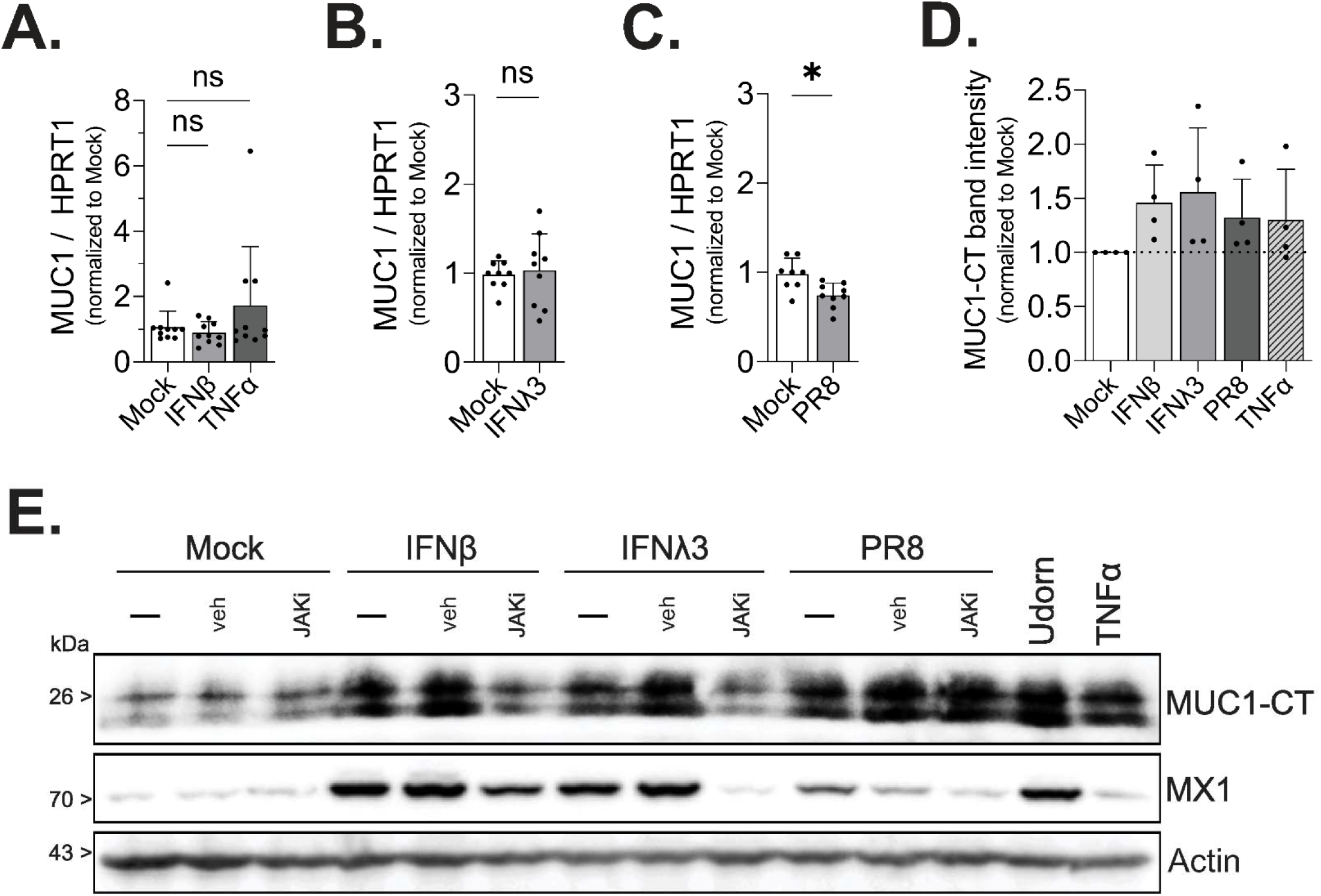
Cell-associated MUC1 levels are upregulated during IAV infection and interferon treatment. HAE were (A and B) stimulated as indicated or (C) infected with IAV and MUC1 expression quantified by qPCR after 24 hours of treatment. In (D) HAE were stimulated as indicated or infected with PR8 as in (A-C) for 24 hours, protein lysate collected, and MUC1 expression quantified by Western blot for MUC1-CT. MUC1-CT band intensity was analyzed by densitometry relative to actin band intensity. In (E), HAE were stimulated with IFN or IAV as indicated (-), in the presence of JAK inhibitor Ruxolitinib (JAKi), or with DMSO as a vehicle control (veh). After 24 hours, lysate was collected and analyzed by Western blot for MUC1-CT, MX1, or actin. In (A-C) results are representative of three biological replicates, and the densitometry analysis of (D) is representative of four biological replicates, including select lanes in (E), all of which are comprised of at least two unique HAE donors. All experimental results were analyzed by Mann-Whitney U test compared to mock conditions and significant where indicated (* p<0.0332).

In order to visualize which cells were expressing MUC1 after IFN challenge or IAV infection, we fixed cultures either 6 and 24 hours post-IFN treatment or 24 hpi and stained for MUC1 using standard immunohistochemical and *en face* immunofluorescence-based approaches. Surprisingly, despite a lack of protein expression in basal cell populations at baseline and a lack of mRNA upregulation after IFN treatment (**S1 Fig** and **Fig 3A**), we observed MUC1 protein in all HAE component cell types following IFNβ stimulation (**Fig 4A**). Infection of HAE with IAV led to increased expression of MUC1 protein by 24 hpi (**Fig 4B**) in both infected and uninfected cells (**Fig 4C**). In line with this broad protein upregulation, both ciliated and non-ciliated cells had increased MUC1 protein as compared to uninfected baseline conditions (**Fig 4D**). Together, our data show MUC1 protein expression is broadly increased in HAE after IFN exposure and IAV infection.

**Fig 4.**
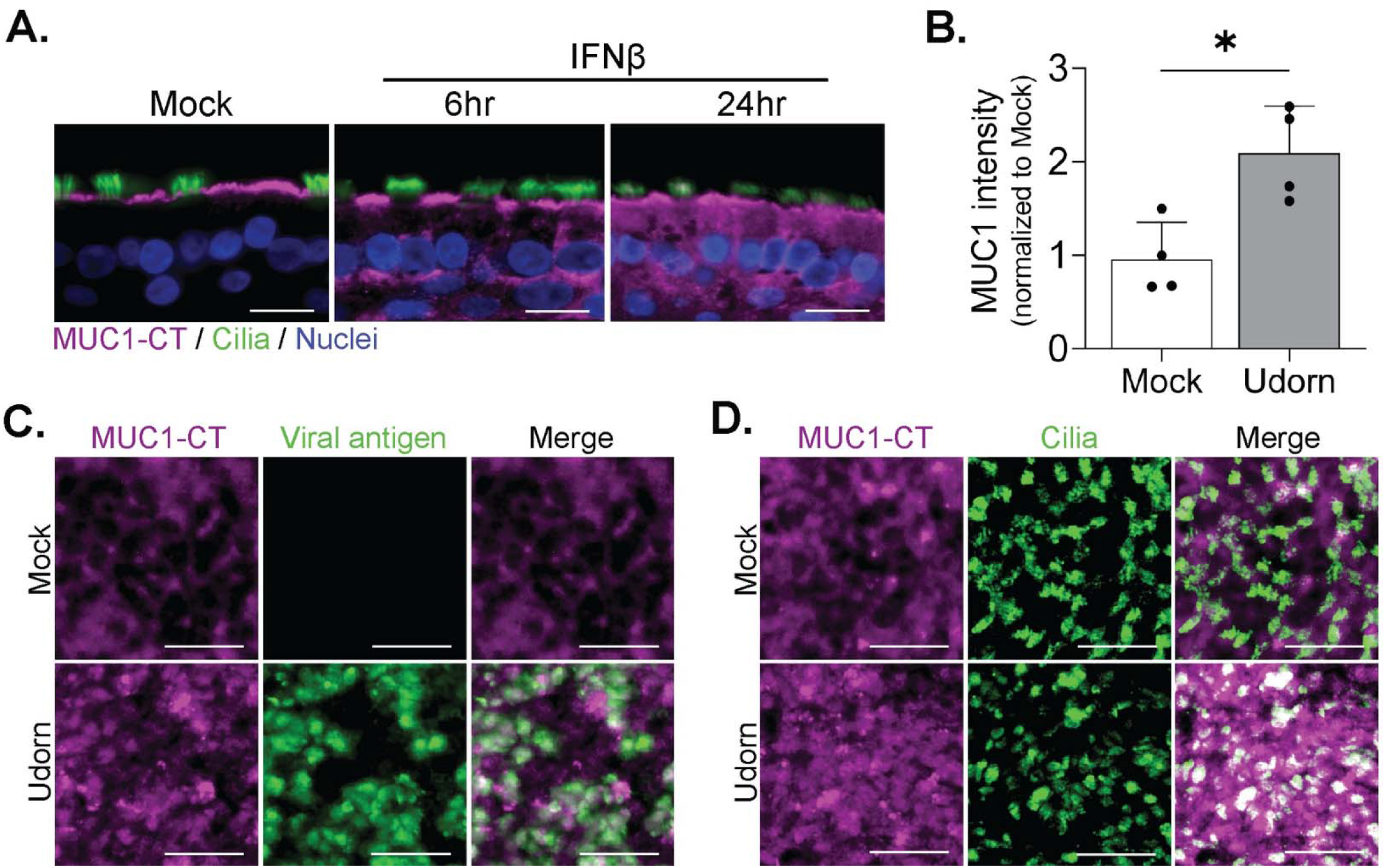
IAV and type I IFN broadly upregulate MUC1 expression across HAE. (A) HAE were stimulated with IFNβ or mock conditions and at indicated time points fixed for immunohistochemical detection of MUC1-CT (purple), acetylated alpha-tubulin (cilia marker; green), and nuclei. HAE were infected with IAV (5 × 10^4^ PFU) and stained *en face* for (B-D) MUC1-CT (purple), (C) viral NP (green), and (D) acetylated alpha-tubulin (cilia marker; green). In (B), mean intensity of MUC1-CT staining was quantified by ImageJ and analyzed by Mann-Whitney U test compared to mock condition, indicating significance (* p<0.0332). Results in (B) are representative of four biological replicates across two unique HAE donors. Scale bars are (A) 20 μm and (C and D) 25 μm.

### Soluble factors secreted by HAE during IAV infection upregulate MUC1 on primary human monocyte-derived macrophages

Beyond epithelial cells, MUC1 is known to be expressed by cells of the hematopoietic lineage [38–40], including macrophages, and this expression can modulate their phagocytic activity [35]. As macrophages play an important role during IAV infection [41,42] and because we observed elevated MUC1 protein during IAV infection and after IFN treatment across HAE component cell types, we next determined the impact of host- and viral-derived factors likely present in epithelial tissue during IAV infection on MUC1 expression in macrophages. Following differentiation with granulocyte-macrophage colony-stimulating factor (GM-CSF; to better achieve alveolar-like macrophages) [43– 45], we stimulated PMD macrophages with Poly I:C (a viral double-stranded RNA mimetic), inflammatory cytokine TNFα, type I interferon (IFNβ), or type III interferon (IFNλ3). Both Poly I:C and IFNβ resulted in a strong upregulation of MUC1 protein similar to the IFNγ (type II IFN) and LPS combination treatment [35] while IFNλ3 and TNFα induced detectible, albeit weak, upregulation (**Fig 5A** and **5B**).

**Fig 5.**
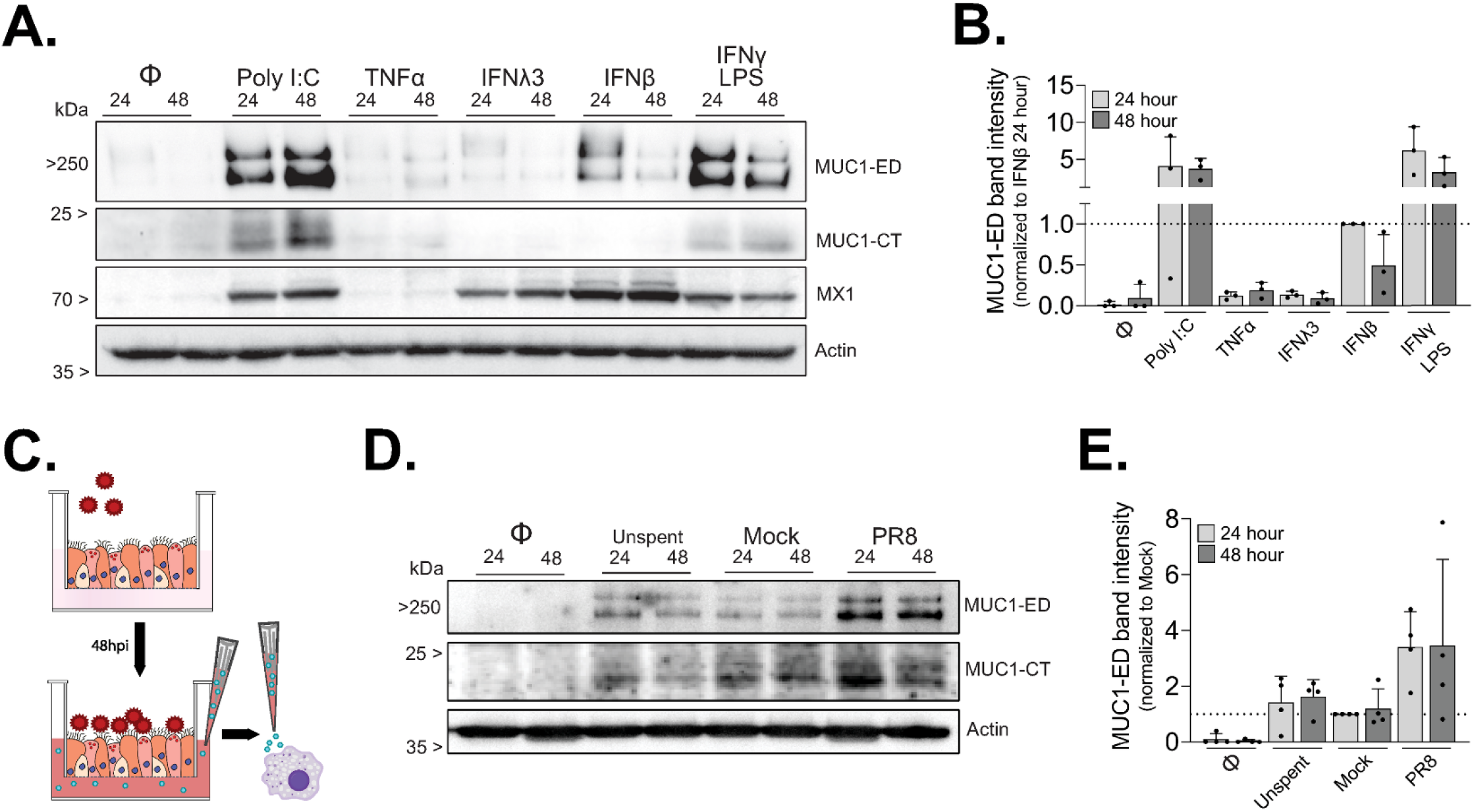
Primary human monocyte-derived macrophages upregulate MUC1 in response to IFN and soluble factors produced from IAV-infected HAE. GM-CSF-derived PMD macrophages were either untreated (Φ) or stimulated as indicated for 24 or 48 hours. (A) Cell lysates were then collected and analyzed by Western blot for MUC1-ED, MUC1-CT, MX1, and actin. (B) Densitometry across conditions reported in (A). Data represent MUC1-ED relative to actin for each sample, normalized to IFNβ / 24hr. (C) Cartoon schematic of experiment conditionsin (D) where PMD macrophages were stimulated with freshly prepared, mock-conditioned, or IAV-infected HAE-conditioned basolateral media before lysate collection and Western blot analysis for MUC1-ED, MUC1-CT, and actin. HAE-conditioned basolateral media used in (D) was collected from independent experimental replicates across unique HAE donors. (E) Densitometry across conditions reported in (D). Data represent MUC1-ED relative to actin for each sample, normalized to Mock / 24hr. Densitometry reported in (B) and (E) represent biological replicates across three and four unique donors, respectively.

To further assess whether MUC1 upregulation was mediated by soluble factors produced in the context of infection, we infected HAE with 5 × 10^4^ PFU of A/Udorn/307/72 and then transferred the virus-free basolateral medium [46] to naïve PMD macrophages (**Fig 5C**). While MUC1 protein was elevated by unspent and mock-conditioned medium, these levels were markedly increased in cultures receiving IAV-conditioned supernatant at both 24 and 48 hours (**Fig 5D** and **5E**). These data indicate that IAV infection of HAE leads to the secretion of soluble factors that have the potential to increase MUC1 levels on multiple cell types during infection *in vivo*.

### Generation of HAE cultures lacking MUC1

Given the ability of IAV to bind MUC1 during infection, and our observed changes in MUC1 protein dynamics in both HAE and PMD macrophages as a consequence of IAV infection, we next sought to determine the impact of MUC1 on IAV replication. We utilized CRISPR/Cas9-mediated genome editing to achieve well-differentiated HAE cultures that were genetically knocked-out (KO) for MUC1. To do so, we cloned a single guide (sg)RNA targeting MUC1 (exon 5; **Fig 6A**), or no known sequence (non-targeting control), into a GFP-expressing lentiviral vector that also encodes the Cas9 nuclease, transduced immortalized human airway epithelial cells (BCi-NS1.1; [47]), and sorted for GFP-positive cells prior to differentiation. Our data demonstrate on-target editing (**Fig 6B**), loss of MUC1 protein (**Fig 6C**), and lack of overt histopathology in differentiated cultures (**Fig 6D**). Compared with control cultures, MCC was significantly reduced in MUC1-depleted cultures (**Fig 6E**); nonetheless, overall, MUC1 was not critical for HAE differentiation or survival, allowing for mechanistic dissection of its role in HAE.

**Fig 6.**
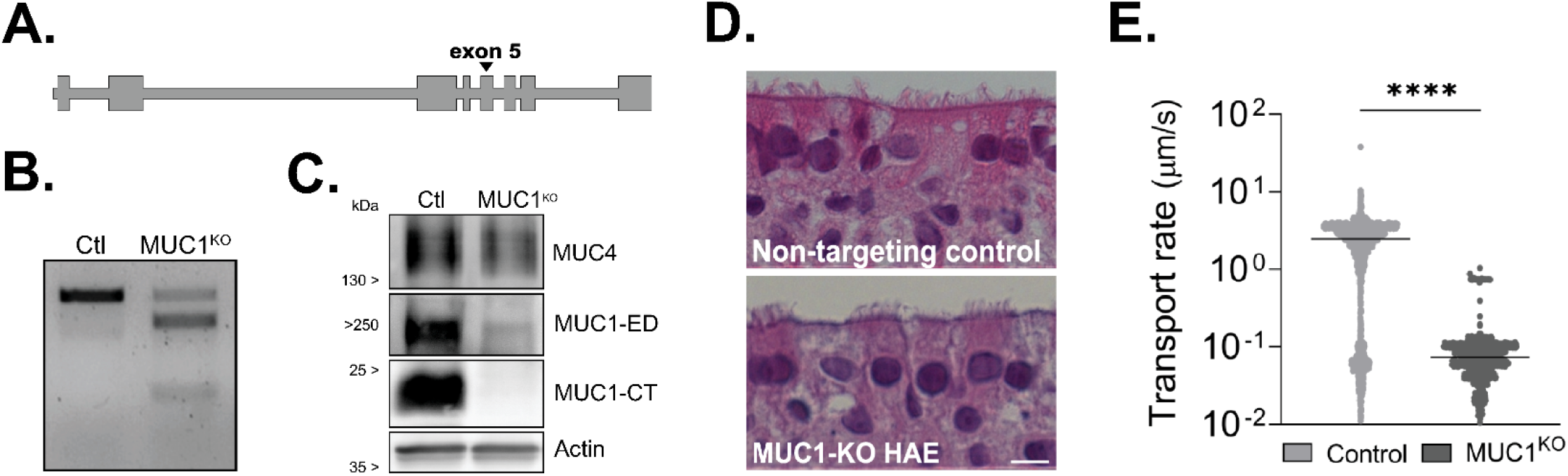
Establishment and characterization of immortalized HAE depleted for MUC1. Immortalized airway epithelial BCi-NS1.1 cells were transduced with CRISPR/Cas9 and sgRNA targeting (A) MUC1 exon 5 for protein depletion (MUC1^KO^) or without predicted targeting site (Ctl / Control). (B) Genomic DNA was extracted and used in a T7 endonuclease I cleavage assay demonstrating editing at the target site. (C) After differentiation, total HAE lysate was collected, separated by PAGE, and blotted for non-targeted tethered mucin MUC4 (extracellular domain), MUC1-ED, MUC1-CT, and actin. (D) H&E stained, histological sections of paraffin embedded cultures show normal ciliated epithelium. Scale bar = 20 μm. (E) Fluorescent microparticles were applied apically to indicated cultures to determine mucociliary transport rate. MCC between culture types was analyzed by Mann-Whitney U test, indicating significance (**** p<0.0001).

### IAV challenge in HAE lacking MUC1 reveals altered infection dynamics

To determine how MUC1 depletion would impact IAV infection dynamics, we inoculated both MUC1 KO and control HAE cultures with 500 PFU A/Udorn/307/72 to allow for multiple rounds of infection and monitored both viral growth kinetics as well as spread throughout the culture by *en face* staining for viral antigen. Viral titers were significantly higher in MUC1 KO cultures compared with control cultures at both 12 and 24 hpi; however, this difference was lost by 48 hpi (**Fig 7A**). These data were consistent with immunostaining results that revealed a limited number of viral antigen-positive cells in control cultures at 12 hpi, while all MUC1 KO cultures had resolvable foci indicative of multicycle replication by this same time point (**Fig 7B**). To assess whether IAV was better able to initiate successful infection of MUC1 KO cultures, we tabulated the number of viral antigen-positive foci on predetermined regions of infected cultures 12 hpi. MUC1 KO cultures had significantly more resolvable foci compared with control HAE cultures (**Fig 7C**). We further expanded this analysis to assess the area of each identified foci, finding that IAV foci were larger in MUC1 KO cultures (**Fig 7D**). In line with these observations, MUC1 KO cultures also had a significantly greater percentage of viral antigen-positive epithelium at 24 hpi (**Fig 7E**). By 48 hpi, both sets of cultures were extensively infected (**Fig 7B** and **Fig 7E**) and the integrity of the apical layer was severely compromised with many regions entirely absent, indicating exhaustive infection in the culture and cytopathic effects (**S4 Fig**).

**Fig 7.**
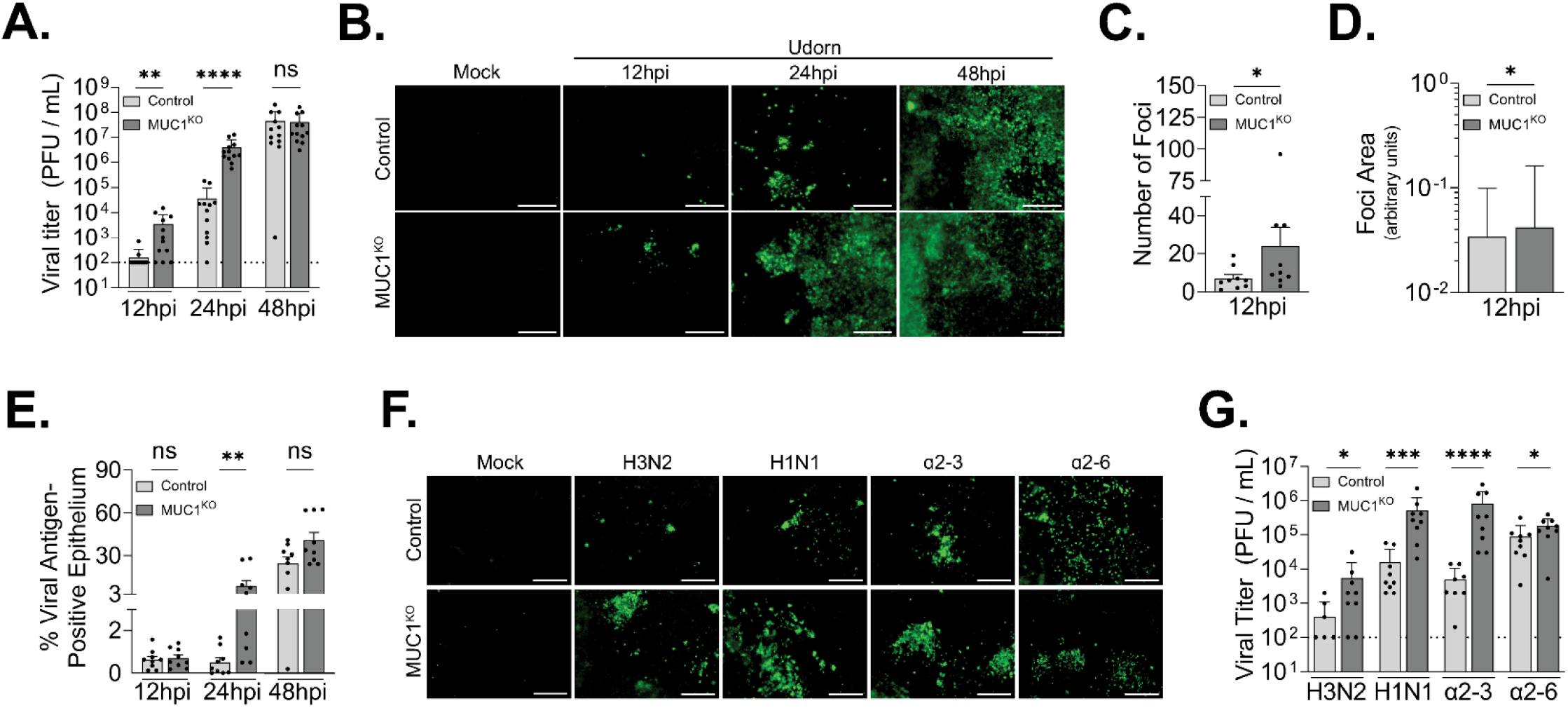
Infection in HAE lacking MUC1 with multiple IAV strains reveals enhanced viral spread. Well-differentiated control or MUC1^KO^ HAE cultures were infected with indicated IAV strains at a low multiplicity of infection (500 PFU). (A-E) At the indicated time points, cultures were washed apically with PBS for (A) viral titer determination, subsequently fixed, and (B) stained *en face* for viral NP antigen. Viral antigen immunofluorescence signal at pre-determined fields of view from HAE was analyzed for the (C) total number of fluorescent foci per individual HAE culture and (D) signal area of contiguous viral antigen (i.e., adjacent infected cells) by culture type. (E) The total viral antigen signal area per culture reported by collection time point. In (F) and (G), cultures were infected with indicated viruses as before and at 24 hpi the apical surface was washed with PBS to (G) determine viral titer, and fixed to be (F) stained for viral NP antigen. Viruses used: H3N2, A/Perth/16/09; H1N1, A/California/04/09; α2-3, A/Udorn/307/72 with HA: L242Q / S244G; α2-6, A/Udorn/307/72(H3N2) with HA: E206D. The results of three experimental replicates in (C), (D), and (E) were determined using ImageJ and all results analyzed by Mann-Whitney U test. Results in (A) and (G) are from four and three experimental replicates, respectively. All data are significant where indicated (* p<0.0332, ** p<0.0021, *** p<0.0002, **** p<0.0001). Scale bars = 100 μm.

As the SA-binding capability of IAV is critical in mediating its endocytic uptake [27], and as we have previously only explored the well-characterized lab strain A/Udorn/307/72, we sought to address whether MUC1’s anti-IAV functionality extends to more recent clinical isolates and A/Udorn/307/72 with altered SA-binding preferences. To address this question, we selected two viruses representing a H3N2 (A/Perth/16/09) and a H1N1 (A/California/04/09) circulating in humans in 2009. Additionally, we created two sets of mutations in the background of A/Udorn/307/72 (capable of binding to both α2-3- and α2-6-linked SA) which lead to enhanced recognition of either α2-3-(HA: L226Q / S228G) or α2-6-linked (HA: E190D) SA [48,49]. MUC1 KO and control HAE were infected as before and both viral titers in the apical compartment as well as the frequency of infected cells were assessed at 24 hpi. As in (**Fig 7B**), all viruses displayed enhanced spread in MUC1 KO cultures compared to control cultures (**Fig 7F**). Furthermore, in congruence with our earlier results, all viruses replicated to higher infectious titers in MUC1 KO cultures (**Fig 7G**) although the magnitude of this difference varied between viruses. Together, these results indicate that in our experimental conditions, MUC1 is not required for initial attachment in HAE and moreover that its loss leads to enhanced viral replication and spread, particularly at early time points (**Fig 7A** and **7G**).

## Discussion

It has been demonstrated that MUC1 plays an important, pathogen-specific, and potentially multifaceted role during respiratory infection [8,14,21,30,35]. MUC1 is an abundant constituent of the PCL where its extracellular domain contributes to airway surface hydration and its cytoplasmic domain has been shown to influence a variety of cellular signaling pathways that modulate the immune response [22,50], cell survival [51,52], and cancer progression [53]. Additionally, MUC1 expression and phosphorylation state depend on external inflammatory stimuli [23,24]. Based on our previous work [33] and that of others [30,35], it is clear that MUC1 plays an important role during IAV infection. However, the nature of this role is poorly understood, and prior research was done in cell culture systems lacking a well-developed glycocalyx or in mice, where mucin orthologs exhibit incomplete homology with their human counterparts. Thus, we sought to explore MUC1-IAV interaction, dynamics of expression, and overall impact on IAV infection in a physiologically-relevant *in vitro* model of human airway epithelium.

Our results support a direct interaction between IAV and endogenous MUC1 during infection in HAE, extending previous findings that demonstrated colocalization of IAV with MUC1 on the surface of A549 cells [30]. Notably, MUC1-ED, the large extracellular domain of MUC1, is capable of dissociating from MUC1-CT through the autocatalytic SEA-module in response to external stimuli [10,54] and it has been previously suggested that this cleavage domain facilitates release of MUC1-ED upon interaction with IAV in the airway lumen [30]. However, we observed a decrease in soluble MUC1-ED after IAV infection in both HAE (**Fig 2**) and A549 cells (**S2 Fig**), suggesting that IAV binding to MUC1-ED does not induce its shedding to a significant degree in systems with endogenous expression with or without a dense glycocalyx. Indeed, we observed an increase in cell-associated MUC1 protein expression in infected cultures (**Fig 3D** and **3E**; **Fig 4B**); thus, a global downregulation of MUC1 cannot account for the loss of soluble MUC1. As conditioned supernatant and culture washes were collected hours after infection in our shedding experiments, it is possible that IAV infection downregulates MUC1-containing vesicles or sheddase expression, resulting in reduced MUC1 release through an indirect mechanism. Alternatively, the near-saturating levels of IAV used in these experiments might reduce MUC1 levels at the cell surface through endocytosis during viral uptake, thereby sequestering it from sheddase activity. As a decreased glycosylation state of MUC1 has been shown to increase its endocytosis [55], this potentially outlines a direct mechanism for IAV glycosidase in reduced surface-expressed MUC1 [56].

Surprisingly, we found that type I and type III IFN can upregulate cell-associated MUC1 protein in HAE (**Fig 3D, 3E**, and **Fig 4A**) despite no significant increases in MUC1 mRNA levels (**Fig 3A and 3B**). These data suggest that MUC1 expression is regulated through a post-transcriptional mechanism under these conditions. IAV upregulation of MUC1 protein in HAE was not exclusively dependent on IFN signaling, indicating multiple soluble factors produced during infection may contribute to elevated MUC1 expression (**Fig 3E**). At least part of this increased expression was due to MUC1 upregulation at the apical surface (**Fig 4C** and **4D**) though broad expression of MUC1 across all HAE component cell types (**Fig 4A and 4D**) after IAV infection and IFN stimulation further indicates that MUC1 expression is nearly ubiquitous across the epithelium. While upregulation at the apical surface likely contributes to barrier function, expression here and in other cells types (e.g., basal cells) may serve alternative roles, potentially suppressing inflammation [50], and/or priming for epithelial repair in response to damage [9,53,57].

As macrophages play a key role during IAV infection [41,42] and previous work demonstrated that macrophages can express MUC1 in response to type II IFN [35], we explored whether IFN produced during IAV infection [36] could induce MUC1 in PMD macrophages. We show here that, in addition to HAE, PMD macrophages upregulate MUC1 following type I and type III IFN stimulation (**Fig 5A** and **5B**). While literature on the human monocyte response to type III IFN is conflicting [58]; human monocyte-derived macrophages and *ex vivo* human macrophages are capable of responding to type III interferon [58–60], which is consistent with our observations across multiple donors. Moreover, these PMD macrophages upregulate MUC1 in response to soluble factors produced by infected HAE (**Fig 5D** and **5E**). These results suggest that sites of infected epithelium might induce MUC1 expression in local macrophages as well as potentially other immune effector cells that have been shown to at least conditionally express MUC1 [38–40]. Interestingly, the banding pattern of MUC1-ED as expressed in PMD macrophages (**Fig 5**) suggests an altered glycosylation state. As the expression [35] and glycosylation state [55] of MUC1 can both independently influence uptake of foreign material in different cellular contexts, further investigation should be undertaken to explore cell-specific impacts of MUC1 expression during IAV infection.

We have also established a MUC1-depleted HAE system through CRISPR/Cas9 technology (**Fig 6**). Others have established similar workflows [61,62] which offer the powerful ability to genetically manipulate otherwise intractable primary human tissue. Our characterization of these immortalized KO cultures reveals robust protein depletion (**Fig 6C**) as well as no gross morphological pathology (**Fig 6D**). Additional functional characterization, however, revealed that MUC1-depleted cultures displayed markedly lower MCC compared with non-targeting control cultures (**Fig 6E**). As the PCL contributes to airway hydration and therefore proper secreted mucus mobility [1–3], it follows that MUC1 depletion could negatively affect this capability. It is also possible that loss of MUC1 alters other factors which impact MCC, such as baseline secreted mucin expression, which were not measured in this study. Future studies on air surface liquid characteristics such as PCL density and/or height, combined with other mucus steady state kinetics (e.g., secreted mucin expression) will better delineate the contribution MUC1 and other tethered mucins make toward overall mucociliary function. The HAE system we utilized here is one of several *in vitro* models that offer the ability to probe the mucosal interface which has been difficult to study in normal 2D tissue culture systems [32].

In our HAE system depleted for MUC1, we found that IAV growth kinetics are increased over control cultures, particularly at 12 and 24 hpi (**Fig 7A** and **7G**). By the earliest time point of 12 hpi, MUC1 depleted cultures had detectable viral titer whereas the majority of control cultures were below the limit of detection (**Fig 7A**). Additionally, not only was the number of foci detectable by *en face* immunofluorescence significantly higher (**Fig 7C**), but there was also clear evidence of multicycle replication visible as early as 12 hpi (the earliest time point investigated) in MUC1 KO cultures compared to control cultures (**Fig 7B** and **7D**). Since IAV can produce new virions as early as 6 hours [63], this implies that there is a significant delay in both the timing and success rate of productive infection initiation in control cultures relative to MUC1-depleted cultures. Consistent with other findings [30], we found that IAV can interact with MUC1, although its absence does not seem to preclude productive IAV uptake. Indeed, as loss of MUC1 enhances viral replication, it is possible that MUC1 may not only be dispensable for initial attachment but in fact may counteract subsequent productive virion adsorption and possibly endocytic entry as well. Importantly, recent work with Muc1 knockout mice has similarly shown that the loss of Muc1 enhances the rate IAV replication, though this did not impact the cumulative viral load [64].

One current model for IAV uptake suggests that virions rely on multivalent interactions with sialylated host proteins and glycolipids to deform local membrane orientation and subsequently trigger endosomal uptake [65,66]. While neuraminidase is normally thought of as a mechanism to avoid virion aggregation and inhibition by secreted mucins [56], recent work has additionally highlighted its importance at this early entry step at or near the host cell membrane [67,68]. In this model, tethered mucins support virion clearance through air-surface liquid hydration and MCC [1,2], but additionally as large constituents of the PCL, also sterically block and, when sialylated, compete with productive virion attachment to membrane-adjacent sialylated attachment sites [1,2]. Our results utilizing wild type Udorn (that binds both α2-3- and α2-6-linked SA) and the mutant Udorn L226Q / S228G and E190D viruses possessing altered SA-binding profiles indicate that MUC1 can antagonize IAV replication regardless of this receptor preference (**Fig 7A and 7G**). Notably, the α2-6-linked SA-binding Udorn mutant (E190D) displayed a much smaller difference in replication between MUC1 KO and control cultures relative to both the α2-3-linked SA-binding mutant (L226Q / S228G) and wild-type Udorn (**Fig 7G**). A preference for α2-3-linked SA might be more inhibited by MUC1 relative to a mixed or α2-6-linked dominant binding profile. However, compared to wild-type Udorn, both SA-binding mutants were attenuated in their replication potential on MUC1 KO cells. Previous work has shown that HA receptor binding preference fitness is also significantly reliant on epistatic balance even between residues outside the receptor binding domain [69] which can confound conclusions about MUC1’s influence on these mutants. Additionally, MUC1 significantly inhibited the replication of both H3N2 and H1N1 clinical isolates (**Fig 7G**). H3N2 and H1N1 viruses have converged in their human receptor preferences for α2-6-linked SA-containing glycoconjugates [70], though more modern drifted H3N2 variants might have continued to diverge in this regard [71]. Both of these clinical isolates display a wider degree of enhanced replication in MUC1 KO relative to control cultures compared to the α2-6-linked SA-binding Udorn mutant. It is also possible that preference for the carbohydrate core in addition to the terminal sialylated moiety further influences the inhibitory function of MUC1. Nonetheless, work on artificial tethered mucin analogs has shown that both sialylated and unsialylated artificial tethered mucins can antagonize productive interactions with gangliosides and delay IAV fusion events, respectively [72]. Together these results suggest that, regardless of receptor binding, MUC1 can significantly antagonize IAV spread in HAE.

Our results are also consistent with the emerging role of MUC1 in response to inflammatory stimuli and we expand on known inflammatory triggers for its expression both in HAE and in PMD macrophages. Indeed, the surprising finding that MUC1 is upregulated beyond the apical layer supports a broader dynamic role during infection at the epithelial surface. Specifically, our data support the model proposed by Kato *et al*. [50] whereby pathogenic insult leads to general inflammation that subsequently upregulates MUC1 expression and immune cell recruitment [64]. This would immediately protect local epithelial cells by acting as a barrier, but further accumulation would help resolve potentially harmful inflammation and simultaneously prime cells for survival and ultimately proliferation to repair local tissue damage following infection.

Additionally, our results demonstrate that MUC1 significantly reduces IAV replication by acting early in infection, consistent with its canonical role as a barrier protecting the airway epithelium. However, instead of the model that suggests MUC1 is acting as a soluble decoy receptor that is dynamically shed in response to viral interaction, our work indicates that MUC1 acts as a general barrier to productive endocytic uptake. As we only investigate the earliest steps in IAV infection of HAE, future studies should aim to investigate how viral-mediated upregulation of MUC1 might impact subsequent spread and immune response to an established infection.

## Materials and Methods

### Human airway epithelial cultures

Human airway tracheobronchial epithelial cells isolated from airway specimens from donors without underlying lung disease were provided by Lonza, Inc. Primary cells derived from single patient sources were expanded on plastic and subsequently plated (3.3×10^4^ cells / well) on rat-tail collagen type 1-coated permeable transwell membrane supports (6.5 mm, #3470; Corning, Inc.). The immortalized HAE line BCi-NS1.1 was kindly provided by Drs. Matthew Walters and Weill Cornell [47].

Both primary and BCi-NS1.1 airway epithelial cells were first expanded in Pneumacult-Ex or Pneumacult-Ex Plus medium (#05008, #05040; StemCell Technologies), and differentiated in Pneumacult-ALI medium (#05001; StemCell Technologies) or custom ALI media (Spirovation, UNC Marsico Lung Institute) with provision of an air-liquid interface for approximately 6 weeks to form polarized cultures that resemble *in vivo* pseudostratified mucociliary epithelium. All cell cultures were maintained at 37°C with 5% CO_2_. All experiments utilized at least two unique donors with figure data indicative of biological replicates.

### Primary human macrophage cultures

Peripheral blood was collected from healthy volunteers, and mononuclear cells were separated by Ficoll-Hypaque density gradient centrifugation. Monocytes were isolated by adherence to plastic and then cultured for one week in X-VIVO 15 serum-free medium (Lonza, Inc.) and 20 ng / mL recombinant human GM-CSF (300-03, Peprotech). Media containing growth factors was replenished 4 days after initial culture. Prior to stimulation, growth factor-containing media was removed and replaced with X-VIVO 15 media supplemented with 5% fetal bovine serum (Genclone). For HAE media stimulations, ALI media was added to standard stimulation media (comprising additional 25% volume) at 24 and 48 hours prior to lysate collection. All studies on human monocyte-derived macrophages were approved by the University of Maryland Institutional Review Board and formal written consent was obtained where necessary.

### MUC1 immunoprecipitation

MUC1 antibodies (B27.29 and 115D8, gifts from Fujirebio Diagnostics Inc.) were conjugated to aldehyde / sulfate latex beads (Invitrogen). Following incubation with anti-MUC1 antibody, beads were incubated with 1 M glycine and 0.5% BSA to coat any remaining exposed area and prevent non-specific binding of protein during immunoprecipitation. HAE apical secretions were pre-treated with 0.1% triton-X before mixing with anti-MUC1-conjugated beads. Following overnight incubation at 4°C, the beads were washed twice with PBS and resuspended in 6 M urea and SDS-PAGE containing reducing agent. HAE secretions (not mixed with beads) were also resuspended in urea / SDS-PAGE buffer as a control. Samples were then vortexed, boiled, and loaded into a 1% agarose gel for electrophoresis and subsequent transfer to nitrocellulose membranes. Membranes were blocked with 5% milk / Tris-buffered saline and 0.1% (v/v) Tween 20 (TBS-T) before incubating with primary antibodies (anti-MUC1 (B2729; 1:2000), recombinant H3-Fc (a gift from Dr. Wendy Barclay; 1:1000), and anti-MUC16 (OC126, Cell Marque; 1:2000)). Recombinant hemagglutinin proteins were generated by infection of insect cells with a recombinant baculovirus expressing the protein as previously described [73]. Membranes overlaid with rH3-Fc were subsequently probed with biotin-SP-conjugated AffiniPure goat anti-human IgG (Jackson ImmunoResearch; 1:2000). Immunodetection was performed using infrared dye-labeled secondary antibodies (IRDye 800CW Biosciences; each at 1:10,000) and visualized using a Li-Cor Odyssey Infrared Imaging System according to the manufacturer’s protocol.

### ELISA

Soluble MUC1 was quantified by ELISA (EHMUC1, Invitrogen) according to the manufacturer’s protocol. To collect HAE samples prior to analysis, 50 μL phosphate-buffered saline (PBS) was applied to the apical chamber and incubated for 30 minutes at 37°C. Prior to experimentation in A549 adenocarcinoma human alveolar basal epithelial cells, growth media (high-glucose DMEM (11-965-092, Gibco) supplemented with 10% fetal bovine serum (Genclone) was replaced with serum-free DMEM. HAE culture washes and A549 culture supernatants were stored at -80°C prior to analysis. Total soluble MUC1 was calculated based on concentration determined by ELISA and total volume collected.

### Influenza virus

The reverse genetics systems for A/Puerto Rico/8/1934 (H1N1; PR8), A/Udorn/307/72 (H3N2; Udorn), and A/Perth/16/09 (H3N2) [74], were generous gifts from Drs. Adolfo Garcia-Sastre, Robert Lamb, and Jesse Bloom, respectively. Live A/California/04/09 (H1N1) virus was purchased from BEI Resources (NR-13658). To produce Udorn viruses with altered SA-binding, HA mutations E206D (E190D; H1 numbering) and L242Q / S244G (L226Q / S228G; H1 numbering) were introduced into the A/Udorn/307/72 reverse genetics system. Specifically, E206D was achieved through GAA to GAU transversion which is predicted to be codon optimized for both canines and humans. L242Q was achieved through CTG to CAA double mutation rather than the single and human codon preferred CAG transversion as a greater barrier to reversion mutation. S244G was achieved through AGT to GGA double mutation to avoid GC/CG dinucleotide bias and avoid codon bias in both canine and human hosts. For viruses derived directly from reverse genetics systems, infectious virus stocks were produced by plasmid transfection in 293T cells and subsequent co-culture with Madin-Darby Canine Kidney (MDCK) cells [75]. Afterwards, all viruses were amplified by passage in MDCK cells (MOI of 0.01) in the presence of 1.5 μg / mL TPCK trypsin. Virus from clarified supernatant was concentrated and purified through 20% sucrose on a 50% sucrose cushion and final viral titer was determined by standard plaque assay on MDCK cells. Briefly, confluent monolayers of MDCK cells in 12 well plates were washed with PBS prior to addition of 100 μL of viral inoculum diluted in serum free DMEM. This was incubated with periodic agitation for one hour at 37°C before being aspirated and replaced with 0.8% molten agar in DMEM/F-12 (Gibco) and 1.5 μg / mL TPCK trypsin. After agar solidification, plates were incubated at 37°C for 72 hours prior to plaque counting.

For infection in unmodified HAE, cultures were washed with PBS for 15 minutes at 37°C to remove apical secretions and supplied with fresh basolateral medium prior to inoculation with sucrose-purified virus diluted in PBS to a final volume of 50 μL. Inoculum was applied to the apical surface of HAE for 2 hours at 37°C. Following incubation, viral inocula were removed, and cultures were washed with PBS for 10 minutes to remove unbound virus. To better mimic natural infection for time course infections in CRISPR/Cas9-modified, BCi-NS1.1-derived HAE, cultures were washed with PBS for 30 minutes at 37°C then incubated for 7 days to allow recovery of the secreted mucus layer prior to inoculation. In these experiments, sucrose-purified viruses were diluted to 500 PFU in 10 μL PBS and inocula were not removed. For all experiments, progeny virus was harvested at indicated times by performing apical washes with 50 μL of PBS for 30 min at 37°C and stored at - 80°C prior to analysis. To measure cytotoxicity, LDH in apical washes was measured with CytoTox 96 (G1780; Promega) according to manufacturer’s instructions.

### qRT-PCR

RNA was extracted using a RNeasy Mini Kit (Qiagen) according to the manufacturer’s instructions. cDNA was prepared separately with SuperScript III (Invitrogen) per manufacturer’s random hexamer protocol. For qPCR, reactions were carried out using LightCycler 480 SYBR Green I mastermix (Roche) and a LightCycler 480 Instrument II (Roche) at the manufacturer recommended settings. Primer sequences, if available, are listed below:

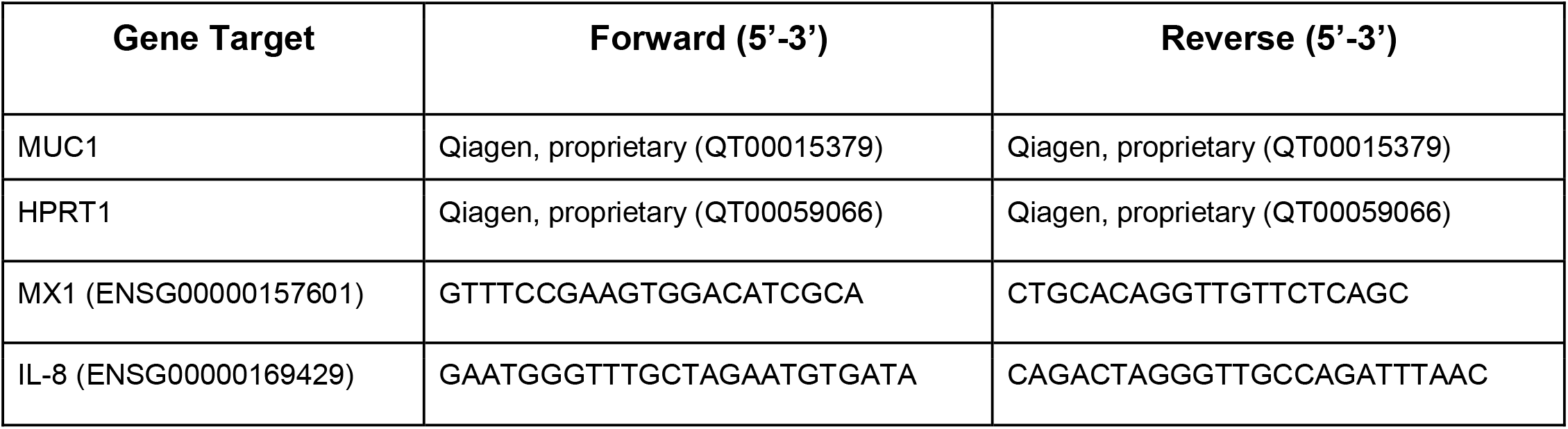

### Western blot

Protein lysate was collected with RIPA buffer (VWR Life Science) supplemented with 2X protease inhibitors (Pierce, Thermo Scientific). Protein concentration was quantified by BCA assay (Pierce, Thermo Scientific), loaded equivalently in each lane (ranging from 4-20 μg between experiments) and run on a 4-20% Tris-Glycine gel (Novex, Invitrogen) under reducing conditions. Protein was transferred to a PVDF membrane (GE Healthcare) and blocked with 5% (w/v) fat free milk protein in TBS-T at room temperature. Unconjugated primary antibody incubations were done in the presence of blocking protein and TBS-T overnight at 4°C. Antibody details are as follows: MUC1-CT (CT2, Invitrogen, 1:5,000); MUC1-ED (B27.29, a gift from Fujirebio Diagnostics Inc., 2.04 μg / mL); MUC4 (1G8, Santa Cruz, 1:5,000); and MX1 (N2C2, GeneTex, 1:5,000). After washing in TBS-T, membranes were probed with secondary antibodies for one hour at room temperature in blocking buffer. Specifically, anti-mouse IgGk-HRP (sc-516102, Santa Cruz, 1:10,000), anti-Armenian hamster IgG-HRP (PA1-32045, Invitrogen, 1:10,000), and anti-rabbit-HRP (32460, Invitrogen, 1:10,000) were used to image MUC1-ED and MUC4, MUC1-CT, and MX1, respectively. Actin was detected using a HRP-conjugated primary antibody (AC-15, A3854, Sigma-Aldrich, 1:35,000) for one hour at room temperature in blocking buffer with rocking. Imaging was performed with chemiluminescent SuperSignal Dura or Femto reagent (Thermo Scientific) on an iBright 1500 (Thermo Fisher). Densitometry in **Fig 3** and **Fig 5** was performed using ImageJ analysis of select band intensity (MUC1-CT or MUC1-ED where indicated) relative to same-sample actin band intensity. Within an experimental replicate, results of individual samples were then normalized to the samples indicated (represented by normalization value of 1.0). MUC1-CT antibodies were used preferentially for Western blot densitometry in HAE due to the smaller size and lack of glycosylation on this part of the mucin. MUC1-ED antibodies were used for densitometry analysis of MUC1 in PMD macrophage experiments as MUC1 expression is significantly lower than that of HAE. The MUC1-ED directed antibody detects a repeated epitope in the VNTR region of MUC1 that enhances sensitivity.

### Cell culture and cell culture treatments

MDCK cells were a generous gift from Dr. Wendy Barclay. They were maintained at 37°C and 5% CO_2_ in high-glucose DMEM (11-965-092, Gibco), supplemented with 10% fetal bovine serum (Genclone) and passaged at 100% confluence with 0.25% trypsin-EDTA (25200-072, Gibco). HEK293T and A549 cells were both purchased through ATCC (#CRL-11268 and #CCL-185). Both HEK293T and A549 cell lines were maintained at 37°C and 5% CO_2_ in high-glucose DMEM (11-965-092, Gibco), supplemented with 10% fetal bovine serum (Genclone) and passaged at 80-90% confluence with 0.05% trypsin-EDTA (25300-062, Gibco). All cell lines were routinely tested for the presence of mycoplasma. Unless specified elsewhere, recombinant human IFNβ (1 nM, 11415-1, PBL Assay Science), IFNλ3 (10 nM, 11730-1, PBL Assay Science), Ruxolitinib (2 μM, S1378, SelleckChem), and DMSO (ATCC) were applied to cell culture media or to both the apical and basolateral chamber of HAE cultures. TNFα (20 ng / mL, 210-TA, R&D Systems) was applied apically to HAE cultures. For experiments with primary macrophage cultures, IFNβ (1 nM), IFNλ3 (10 nM), TNFα (20 ng / mL), low molecular weight Poly(I:C) (10 μg / mL, k-picw, InvivoGen), and LPS (100 ng / mL *E. coli* K12, k-eklps, Invivogen) with IFNγ (20 ng / mL, R&D Systems) were supplemented into X-VIVO 15 media and 5% fetal bovine serum at indicated time points prior to lysis.

### Histology, immunohistochemistry (IHC), and immunofluorescence (IF) microscopy

HAE cultures were fixed in 4% paraformaldehyde overnight prior to paraffin embedding and sectioning at either the Marsico Lung Institute Histology Core (Chapel Hill, NC) or the New York University Experimental Pathology Research Laboratory (New York, NY). Five micron-thick sections on slides were deparaffinized with xylene and rehydrated through gradient ethanol washes into distilled water. For heat antigen retrieval, citrate buffer (2.94 g / L), pH 6.0, with 0.05% Tween-20 was heated to 98°C to boil deparaffinized slides for 15 minutes. After cooling the slides and washing in water, slides were blocked with 3% BSA in PBS supplemented with 1 mM CaCl_2_ and 1 mM MgCl_2_ (PBS++). Primary antibodies were diluted in 1% bovine serum albumin / PBS++ and incubated with the sample overnight at room temperature. Slides were then washed with PBS++ and secondary antibodies (also diluted in 1% bovine serum albumin / PBS++) added for one hour at room temperature. Slides were then stained with 4’,6-Diamidino-2-Phenylindole, Dihydrochloride (DAPI, Invitrogen) or Hoechst 33342 Solution (Thermo Scientific), washed a final time with PBS++, and coverslips mounted with Vectashield antifade mounting solution (H-1000, Vector Laboratories). To prepare HAE cultures for *en face* IF staining, cultures were fixed in 4% paraformaldehyde for 20 minutes at room temperature, permeabilized with 2.5% Triton X-100, and blocked with 3% BSA in PBS++. The IF antibody staining procedure was the same for IHC. For the IF protocols above, antibodies for IHC were as follows: acetylated alpha tubulin (cilia marker, clone 6-11B-1, ab24610, Abcam, 1:2000); MUC1-CT (clone MH1, CT2, MA5-11202, 1:150); anti-Armenian hamster IgG AlexaFluor-647 (ab173004, Abcam, 1:500); MUC16 (clone X325, ab10033, Abcam, 1:1000); MUC4 (clone 1G8, sc-33654, Santa Cruz, 1:100); and anti-mouse IgG AlexaFluor-488 (Invitrogen, 1:500). For IF, antibodies were as follows: acetylated alpha tubulin (clone 6-11B-1, ab24610, Abcam, 1:2000) and anti-mouse IgG2b (clone 7E10G10, ab170192, Abcam, 1:500); MUC1-CT (clone MH1, CT2, MA5-11202, 1:50) and anti-Armenian hamster IgG AlexaFluor-647 (ab173004, Abcam, 1:500); influenza virus NP (clones A1 and A3, MAB8251, Sigma-Aldrich, 1:100) and anti-mouse IgG AlexaFluor-488 (Invitrogen, 1:500). MUC1-CT was utilized to accommodate the use of other mouse antibodies in concurrent staining. Images were acquired with a Zeiss Axio Observer 3 Inverted fluorescence microscope equipped with a Zeiss Axiocam 503 mono camera and Zen imaging software.

For transmission electron microscopy detection of virus and MUC1, two protocols were used. In the first, HAE cultures were washed and 4.7 × 10^6^ PFU sucrose-purified A/Udorn/307/72 was allowed to adsorb for one hour at 37°C followed by transfer of the cultures to 4°C for all subsequent steps up to fixation (**Fig 1E**). In the second, HAE cultures were washed and 5 × 10^5^ PFU dialyzed, sucrose-purified A/Udorn/307/72 was allowed to adsorb for 2 hours at 4°C (**Fig 1C, 1D, and 1F**). Virus inoculum was removed and cultures were blocked with 10% (v/v) normal donkey serum for one hour. Anti-MUC1-ED B27.29 (2.04 ug / mL) and anti-Hong Kong/68 goat antiserum (NR-3118, BEI Resources) was added in the presence of blocking serum overnight. Cultures were washed with PBS++ to remove primary antibodies before addition of 18 nm-gold conjugated anti-mouse (1:10, Jackson ImmunoResearch Laboratories, Inc.) and 6 nm-gold conjugated anti-goat (1:20, Jackson ImmunoResearch Laboratories, Inc.) in blocking solution for one hour. Secondary antibodies were removed, cultures washed and subsequently fixed in 2% glutaraldehyde in 0.1M cacodylate buffer for one hour at room temperature. Following a further washing step in 0.1 M cacodylate buffer, a secondary fixation step using 1% OsO4 and 1% K_3_Fe(CN)_6_ in 0.1M cacodylate buffer was performed for one hour. A final wash of 0.1M cacodylate buffer was performed before post-fixation treatment with 2% uranyl acetate solution in dsH_2_O for one hour. Cultures were then dehydrated in increasing concentrations of ethanol. Finally, cultures were infiltrated with 100% propylene oxide and subsequently increasing ratios of Spurr’s Resin up to the final embedding step of 100% Spurr’s Resin. Cultures were then imaged at 80kV on the Hitachi HT7700 Transmission Electron Microscope at the Laboratory for Biological Ultrastructure at the University of Maryland.

### Mucociliary clearance

HAE mucus was allowed to accumulate for one week prior to the application of 5 μl of 2 μm red-fluorescent (Cy3) polystyrene microspheres (1:50 dilution, Sigma-Aldrich) to the apical chamber of the transwell. Cultures were allowed to equilibrate for 24 hours after which the HAE cultures were imaged. For each culture, videos of three regions were recorded at 10 X magnification. Images were collected at a frame rate of 0.5 Hz for 60 seconds on the plane of the mucus gel. Since the secreted mucus tends to accumulate at the edges of the transwells, images were taken centrally to avoid areas of thick mucus. The microsphere tracking data analysis is based on an image processing algorithm that was custom written in MATLAB [76]. Briefly, the analysis software computes the xy-plane trajectories of each fluorescent microsphere in each frame. Using the first and last position obtained from trajectory data, displacement of microspheres was computed, and transport rate was calculated by dividing the displacement by total time elapsed. Microspheres with transport rates of less than 0.01 μm / s (less than 0.01% of microspheres) were considered immobile and removed from the data set.

### CRISPR/Cas9-mediated knockout of specific mucin glycoproteins in HAE

To select regions for CRISPR/Cas9-mediated knockout, MUC1 (ENSG00000185499) was analyzed using Ensembl [25]. Guide RNA sites were selected based on favorable targeting, Doench, and Xu scores. Putative guides were ordered from IDT with flanking restriction sites for cloning into the plentiCRISPRv2 backbone [77] with eGFP replacing puromycin selection. The final guide targets region 155,187,791 – 155,187,813 on chromosome 1 with WTSI Genome Editing ID of 915343298. To generate negative control gRNA sequences with no matching sequence in the genome (non-targeting control), we generated 10,000 random sequences of 20 nucleotides and analyzed these candidates using BLAST [78] to characterize their match percent identity against the hg19 reference genome. We chose the gRNAs sequence with the least match percent identity i.e. lowest probability of a sequence match in the genome, as the negative controls. To improve the confidence of our hits, we repeated this process at a wide range of coverage thresholds (99-70%) and chose top-ranked candidates consistently ranked among top ones (average rank). We used the following control sequence, which was validated as described above: 5’ CGA CTA CCA GAG CTA ACT CA 3’.

Lentiviral stocks were generated by co-transfection of 1 μg plentiCRISPRv2 (a gift from Dr. Feng Zhang (Addgene plasmid #52961; http://n2t.net/addgene:52961; RRID:Addgene_52961)), 0.2 μg pCMV-VSV-G (a gift from Dr. Bob Weinberg (Addgene plasmid #8454; http://n2t.net/addgene:8454; RRID:Addgene_8454)) [79], and 0.7 μg psPAX2 (a gift from Dr. Didier Trono (Addgene plasmid #12260; http://n2t.net/addgene:12260; RRID:Addgene_12260)) into HEK293T cells with X-tremeGENE HP (Roche) in OptiMEM (Invitrogen) per manufacturer’s protocol. Lentivirus-laden supernatant was collected and replaced at 24 hour intervals up to 72 hours, pooled, and filtered to remove viable cells and debris.

For target cell transduction, lentivirus-containing supernatant was applied to BCi-NS1.1 (kindly provided by Drs. Matthew Walters and Ronald Crystal; maintained as HAE above, [47]) at 40-60% confluence with a final concentration of 20 mM HEPES (Gibco) and 4 μg / mL Polybrene (American Bio). Cells were then centrifuged (1,000 *g* for one hour at 37°C) and incubated at 37°C overnight. The inoculum was removed and replaced with fresh growth media. At 60-80% confluence cells were passaged and expanded prior to being sorted for eGFP expression compared to untransduced control cells. Sorted transduced cells were frozen down for later use or subjected to EnGen mutation detection kit (New England BioLabs) for on-target gene editing confirmation. Upon thawing, transduced cells were expanded once before transfer to collagen-coated membranes as with primary HAE. Target protein depletion in mature, differentiated cultures was confirmed by Western blot. Select cultures were fixed in 4% paraformaldehyde, then paraffin-embedded, sectioned, and stained with hematoxylin and eosin (H&E) at the New York University Experimental Pathology Research Laboratory, and subsequently imaged on a Nikon eclipse microscope at the University of Maryland Imaging Core.

### Software Used and Statistical Analysis

ImageJ was used to quantify fluorescence intensity in IF experiments and band intensity of indicated Western blot developments. Statistical analyses were performed using native GraphPad Prism 8 software.

## Acknowledgments

We thank Drs. Adolfo Garcia-Sastre (Icahn School of Medicine at Mount Sinai), Robert Lamb (Northwestern University), Wendy Barclay (Imperial College London), Jesse Bloom (University of Washington and Howard Hughes Medical Institute), and Matthew Walters and Ronald Crystal (Weill Cornell Medicine) for generously sharing reagents. We also thank the Addgene depositors Drs. Feng Zhang, Bob Weinberg, and Didier Trono for their contributions in making reagents broadly accessible. We acknowledge Dr. Jacques Augenstreich for resources and training in primary human macrophage culture. We are also grateful to the directors and teams of the University of Maryland Laboratory for Biological Ultrastructure, Imaging Core, and Genomics Core as well as the New York University Experimental Pathology Research Laboratory and the Marsico Lung Institute Tissue Culture and Histology Cores.

The following reagents were obtained through BEI Resources, NIAID, NIH: Polyclonal Anti-Influenza Virus H3 Hemagglutinin (HA), A/Hong Kong/1/1968 (H3N2), (antiserum, Goat), NR-3118. Influenza A Virus, A/California/04/2009 (H1N1)pdm09, Cell Isolate (Produced in Cells), NR-13658.

The authors declare that they have no competing interests.

## Authors’ contributions

MAS designed the project. EI and MAS wrote the manuscript and designed the experiments. EI, KG, DS, TBG, KH, MK, and MP performed the experimental work. Specifically: EI rescued, propagated, and concentrated virus stocks, generated transmission electron microscopy samples, generated MUC1 depleted HAE cultures and controls, performed *en face* staining of HAE cultures, and analyzed PMD macrophage lysates by western blot; KG performed and analyzed experiments related to MUC1 expression in HAE cultures; DS performed MCC microscopy and analysis; TBG processed and analyzed transmission electron microscopy samples; KH isolated, differentiated, and stimulated the PMD macrophages; MK performed the MUC1 immunoprecipitation from HAE apical secretions and overlay experiments; SS developed a tool for control guide RNA design; MP and EI performed MUC1 quantitation experiments; GAD and MK contributed reagents and expertise. All authors read and approved the final manuscript.

## Supporting information captions

**S1 Fig. The HAE system recapitulates airway epithelial morphology and tethered mucin expression**. Immunohistochemistry of primary HAE cultures detecting the extracellular domains of tethered mucins MUC1, MUC4, and MUC16. MUC4 and MUC16 stains represent immature glycosylation forms while mature proteins localize to the extracellular apical lumen. Scale bar = 20 μm.

**S2 Fig. MUC1-expressing cells that lack a robust glycocalyx do not shed MUC1 after IAV infection**. (A) A549 cells were stained for the extracellular domain of MUC1 and nuclei. Scale bar = 50 μm. In (B), A549 cells were infected with IAV as indicated and MUC1 in the cell culture supernatants 24 hours post-infection was quantified by ELISA. Results from four experimental replicates were analyzed by Mann-Whitney U test compared to mock condition (ns = not significant; ** p<0.0021).

**S3 Fig. Expression of canonical interferon-stimulated genes and inflammatory chemokines in HAE following cytokine stimulation**. HAE were stimulated with recombinant human IFNβ, TNFα, or mock conditions for 24 hours. Total RNA was then collected and (A) MX1 and (B) IL-8 expression analyzed by qPCR. In (C) HAE were stimulated with recombinant IFNλ3 for 24 hours prior to RNA collection and qPCR quantification of MX1 as before. Results from three experimental replicates and three unique donors were analyzed by Mann-Whitney U test compared to mock conditions (ns = not significant; ** p<0.0021; **** p<0.0001).

**S4 Fig. Relative cytotoxicity increases substantially at later time points during IAV infection in HAE**. Relative cytotoxicity in MUC1-depleted and control HAE following IAV infection determined by quantification of lactate dehydrogenase levels in apical washes at indicated time points. Results from three experimental replicates were analyzed by Mann-Whitney U test compared to control cultures of the same time point (ns = not significant; **** p<0.0001).

