## Supplementary figures and images for "Membrane-Tethered Mucin 1 is Stimulated by Interferon in Multiple Cell Types and Antagonizes Influenza A Virus Infection in Human Airway Epithelium"

### Supplemental Figures

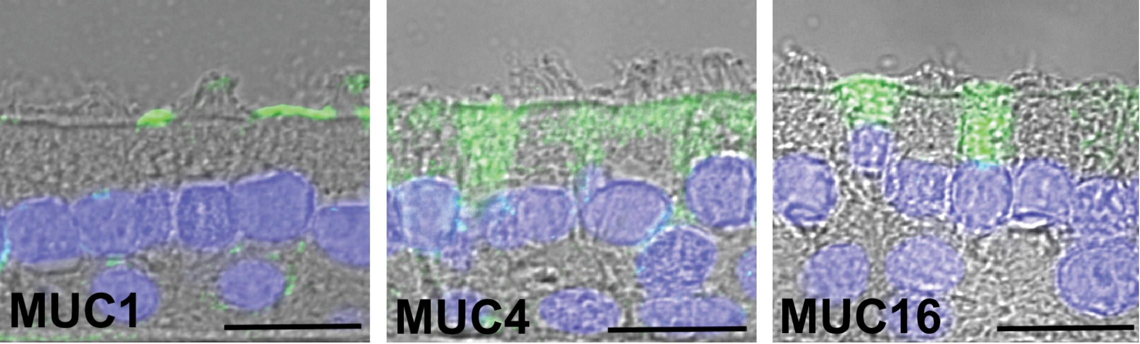

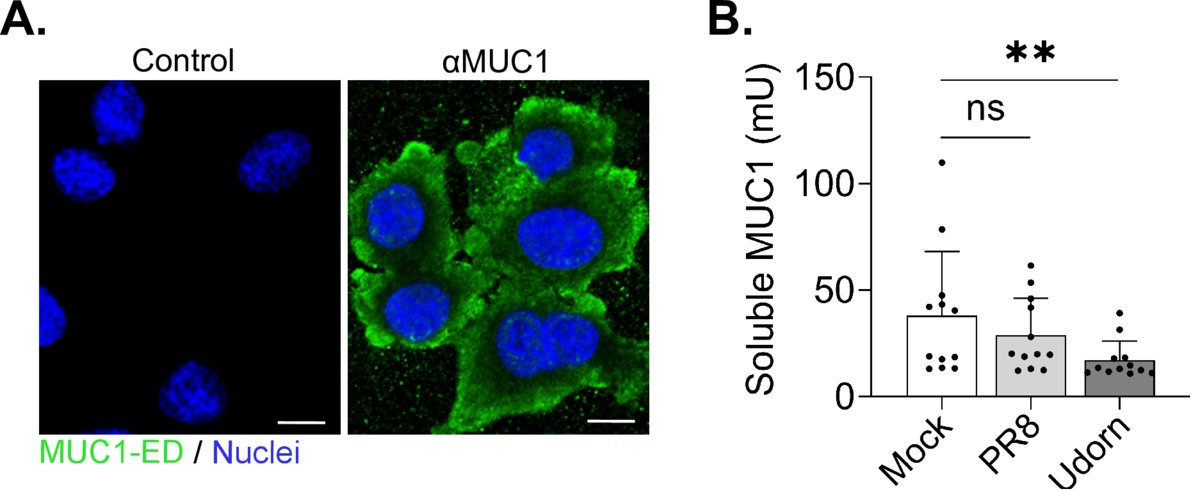

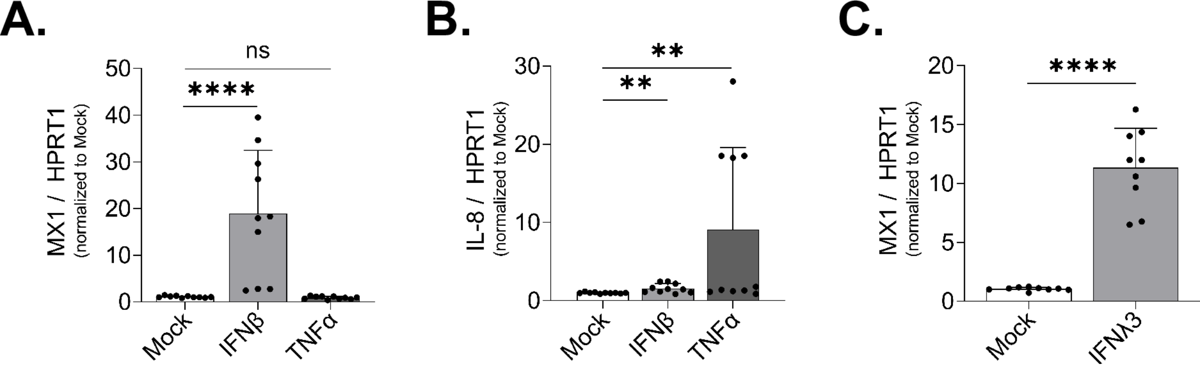

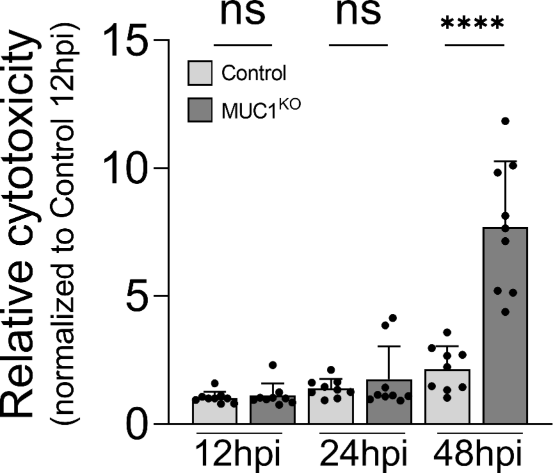
